# Pharmacological CDK4/6 inhibition unravels a p53-induced secretory phenotype in senescent cells

**DOI:** 10.1101/2020.06.05.135715

**Authors:** Boshi Wang, Simone Brandenburg, Alejandra Hernandez-Segura, Thijmen van Vliet, Elisabeth M. Jongbloed, Saskia M Wilting, Naoko Ohtani, Agnes Jager, Marco Demaria

## Abstract

Cellular senescence is a state of stable growth arrest that acts as a tumor suppressive mechanism. Several anti-cancer interventions function partly by inducing malignant cells into senescence. However, because of systemic administration and lack of specificity, anti-cancer treatments are associated with premature senescence of various non-malignant cells. Therapy-induced non-malignant senescent cells can have profound detrimental pro-tumorigenic and pro-disease functions via activation of a pro-inflammatory and NF-κB-mediated secretory phenotype (SASP). Inhibitors of the cyclin-dependent kinases 4/6 (CDK4/6i) has recently shown to have potent cytostatic effects with reduced toxicities. Here, we show that CDK4/6i lead non-malignant cells to a senescent state that lacks the pro-inflammatory and NF-κB-associated SASP. Interestingly, CDK4/6i-induced senescence overexpressed a number of genes encoding for secreted proteins, which we show being dependent on p53 transcriptional activity. CDK4/6i-induced p16^+^ senescent cells with a p53-associated (PASP), but not NF-κB-associated (NASP), secretory phenotype do not exert detrimental and pro-tumorigenic functions, but still retain the capacity to induce paracrine senescence and undergo clearance *in vivo.* Our data suggest that that senescent cells with a PASP but without a NASP may be well-tolerated and may represent a less toxic outcome for cancer interventions.

## Introduction

Cellular senescence is a state of stable growth arrest dependent on the upregulation of cyclin-dependent kinases (CDK) inhibitors p16 and p21, and associated to a complex Senescence-Associated Secretory Phenotype (SASP) (*1*). Senescence induction acts as a potent tumor suppressive mechanism by restraining cancer cell proliferation both via the cell-autonomous cell cycle arrest and by non-cell autonomous induction of paracrine senescence. Moreover, the SASP can potentiate the senescence-associated tumor suppressive functions and regulate senescence turnover by activating immunosurveillance and immune-mediated clearance (*2*). Because of its potent anti-neoplastic function, induction of cellular senescence is a desired outcome of anticancer interventions (*3*), and a well-described phenomenon in pre-clinical and clinical settings that involve the use of genotoxic therapies such as chemo and radiotherapy (*4*). However, lack of specificity and systemic administration of classic anti-cancer interventions are also the basis of several short-and long-term adverse reactions. Increasing evidence suggests that non-malignant cells in various tissues can enter senescence during therapy and promote a state of chronic inflammation and tissue degeneration (*5–8*). Genetic and pharmacological removal of therapy-induced senescent cells is sufficient to alleviate various adverse reactions including fatigue, myelosuppression, cardiomyopathy, bone loss, frailty, cancer progression and relapse (*5–8*). These senescence-associated detrimental effects are mainly mediated by pro-inflammatory SASP cytokines and chemokines, which are normally elevated in cancer patients suffering from a variety of adverse reactions to cancer therapies (*9, 10*). Major regulator of pro-inflammatory SASP genes is NF-κB, a transcription factor mediating the response to persistent DNA damage (*11, 12*). In contrast to DNA damaging events, senescent states activated by direct overexpression of the CDK4/6 inhibitor p16 do not engage a NF-κB-driven and pro-inflammatory SASP (*13–15*). Palbociclib (PD033291) and abemaciclib (LY2835219) are selective CDK4/6 inhibitors (CDK4/6i) approved by the U.S. FDA to treat hormone-sensitive and HER2 negative advanced breast cancer patients. Addition of a CDK4/6 inhibitor to hormonal treatment showed to improve progression free survival in large metastasized breast cancer patient cohorts (*16, 17*) and under investigation as first line treatment for lung cancer patients (*18*). Short-term treatment with CDK4/6i induces a temporary and reversible growth arrest, but long-term treatment promotes a poorly characterized state of cellular senescence (*19, 20*). Here, we set to study the phenotype of non-malignant cells induced to senescence by pharmacological CDK4/6 inhibition, evaluate their non-cell autonomous functions and their toxicity *in vivo*.

## Results

### CDK4/6i treatment induces cellular senescence but not NF-κB-dependent SASP

In accordance with previous studies (*21*), prolonged treatment of human primary fibroblasts BJ with CDK4/6 inhibitors abemaciclib and palbociclib was associated to a progressive loss of RB phosphorylation (fig. S1A) and down-regulation of *E2F2* (fig. S1B) which resulted in cells entering a senescence state characterized by irreversible growth arrest (Fig. 1A and fig. S1, C and D), flattened and enlarged morphology (fig. S1E), activation of the Senescence Associated-β-galactosidase (SA-β-gal) (Fig. 1B and fig. S1F) and induction of the endogenous CDK4/6i p16 (fig. S1G). In contrast to the genotoxic stress inducer agent doxorubicin, treatment with abemaciclib and palbociclib did not trigger a DNA damage response (Fig. 1C). To further compare CDK4/6i and genotoxic stress treatments, we performed RN A-sequencing analysis of cells induced to senescence either by abemaciclib or doxorubicin. Several genes were similarly differentially regulated in the two groups (Fig. 1D), including the ones associated to cell cycle (Fig. 1E), lysosomal-associated pathways (Supplementary Table 1) and a ‘core’ signature of senescence (*22*) (Fig. 1F). In contrast, several pro-inflammatory SASP factors predicted to be NF-κB targets were only induced in the doxorubicin group (Fig. 1G), which followed the lack of a DDR (Fig. 1C) and of NF-κB transcriptional activity in the abemaciclib group (Fig. 1H). qPCR (Fig. 1I), ELISAs (Fig. 1J) and cytokine array assays (fig. S2A) confirmed lack of NF-κB-associated SASP factors in BJ cells induced to senescence by pharmacological CDK4/6 inhibition, either by abemaciclib or palbociclib, whereas a strong induction was observed in cells induced to senescence by doxorubicin and by another chemotherapeutic and genotoxic agent, paclitaxel (*23, 24*). This difference was also observed in cell types other than fibroblasts, including the epithelial cells RPE1 and lung mesenchymal stem cells, which entered a state of senescence with no NF-κB-dependent SASP upon treatment with CDK4/6i (fig. S2, B and C). Finally, when we measured the level of the NF-κB-dependent SASP factors CXCL1 and CCL5 in the plasma of breast cancer patients treated either with paclitaxel or palbocilib, we observed an increase only in patients treated with paclitaxel (Fig. 1K). Altogether, these data demonstrate that treatment with CDK4/6i causes non-malignant cells to enter a state of senescence without NF-κB signaling and of a NF-κB pro-inflammatory secretory program, from now on defined NASP (NF-κB-associated secretory phenotype).

**Figure 1.**
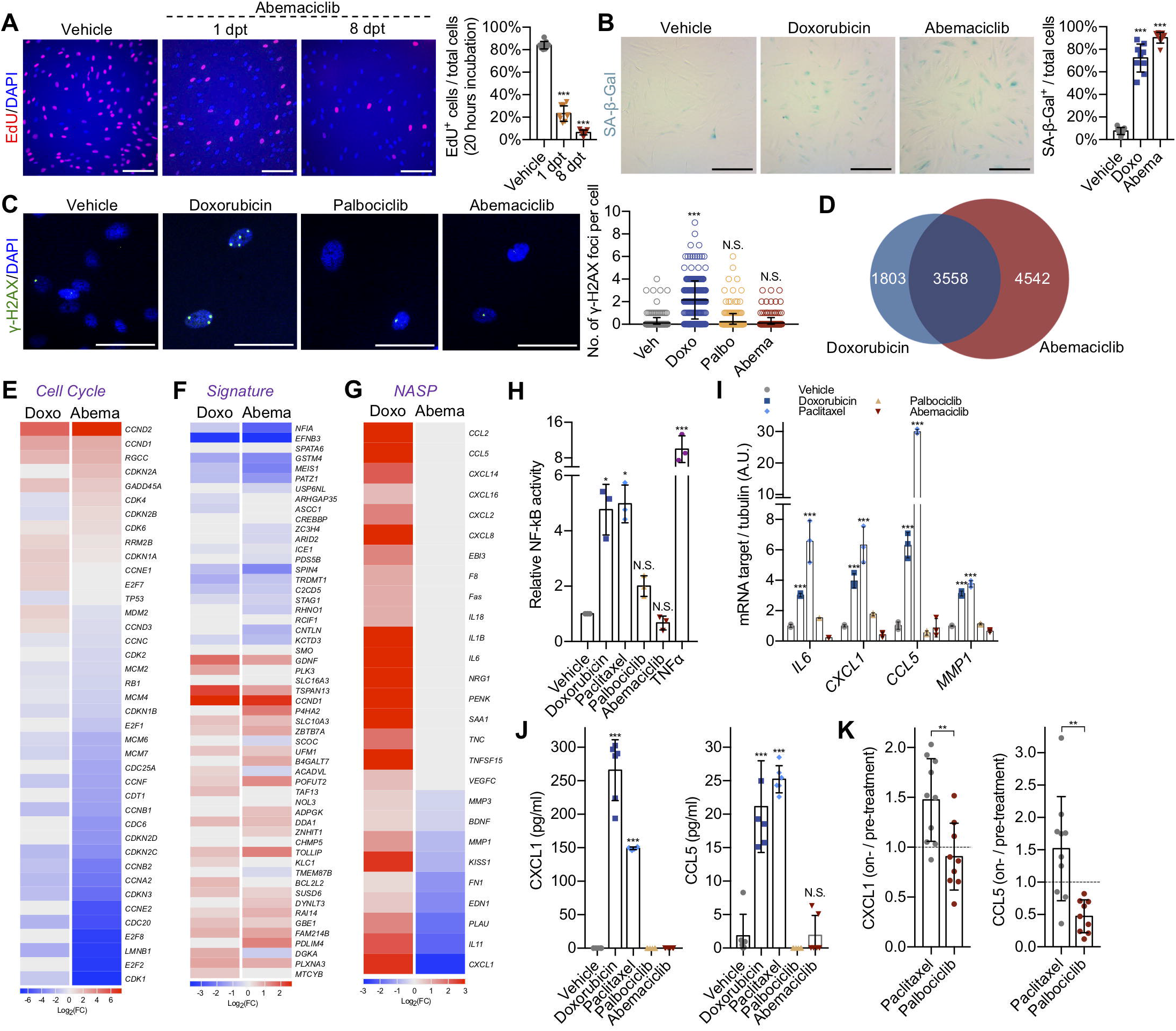
CDK4/6i treatment induces cellular senescence but not NF-κB-dependent SASP. (**A** to **C**) Human fibroblasts (BJ) were treated with vehicle (water or DMSO for 8 times 24 hours) or abemaciclib (1 μM for 1 or 8 times 24 hours) or doxorubicin (250 nM for 24 hours) or paclitaxel (50 nM for 24 hours) or palbociclib (1 μM for 8 times 24 hours). 1 or 8 dpt, cells treated with vehicle or abemaciclib were incubated with EdU for 20 hours and stained and quantified (N=9 samples from 3 independent experiments; scale bar, 150 μm) (**A**). 8 dpt, cells treated with doxorubicin or abemaciclib were stained for SA-β-gal and quantified (N=9 samples from 3 independent experiments; scale bar, 1 mm) (**B**). 8 dpt, cells were stained for γ-H2AX (scale bar, 60 μm) and quantified (vehicle=288 cells, doxorubicin=280 cells, palbociclib=276 cells, abemaciclib=266 cells from 3 independent experiments) (**C**). (**D** to **G**) RNA-sequencing datasets of doxorubicin or abemaciclib treated cells at 8 dpt (N=3 independent samples, sequenced together). Venn plot generated from the datasets (**D**). Heatmap of cell cycle genes (**E**) or the senescence-associated signature (**F**) or the NF-κB-dependent SASP (NASP) (**G**) for doxorubicin- or abemaciclib-treated groups (8 dpt) relative to vehicle-treated group from the RNA-sequencing datasets (p-value<0.01 was regarded as significant). (**H**) BJ cells transduced with a NF-κB reporter were treated with vehicle, doxorubicin, paclitaxel, palbociclib or abemaciclib and luciferase activity measured 8 dpt. TNFα treatment was used as positive control (N=3). (**I**) 8 dpt, qRT-PCR of indicated genes was performed using vehicle-, doxorubicin-, paclitaxel-, palbociclib- or abemaciclib-treated BJ cells (N=3). (**J**) Serum-free conditioned media (CM) was collected from indicated treated BJ cells 8 dpt and expression levels of CXCL1 and CCL5 measured by ELISA (N=6). (**K**) Cell-free plasma was derived from breast cancer patients treated with paclitaxel (N=10) or palbociclib (N=9) and the expression levels of CXCL1 and CCL5 quantified by ELISA. The values represent the ratio between on-treatment and pre-treatment levels for each patient. One-way ANOVA, data are means ±SD (**A, B, C, H**, and **J**). Two-way ANOVA, data are means ±SD (**I**). Unpaired two-tailed *t*-test, data are means ±SD (**K**). *p<0.05, **p<0.01, ***p<0.001, N.S.=not significant. dpt, days post treatment. Doxo, doxorubicin. Abema, abemaciclib.

### CDK4/6i treatment induces a p53-dependent SASP

RNA-sequencing data suggested that despite the absence of SASP pro-inflammatory factors, other previously-described SASP factors were similarly upregulated in both abemaciclib- and doxorubicin-induced senescent cells. Interestingly, several of the SASP factors upregulated in the abemaciclib group were predicted to be p53 target (*25*) (Fig. 2A). Elevated levels of the p53-target and SASP factors *IGFBP3* and *LIF* were observed in BJ, epithelial and mesenchymal stem cells induced to senescence by the CDK4/6i abemaciclib and palbociclib (fig. S3A), and IGFBP3 was elevated in the plasma of breast cancer patients upon treatment with palbociclib (Fig. 2B). To evaluate if CDK4/6i treatment led to elevated p53 transcriptional activity we used a reporter system and Chromatin Immunoprecipitation (ChIP) assays. BJ cells with stable expression of a luciferase reporter construct (p53LUC) responding to p53 direct binding demonstrated gradual activation of p53 upon exposure to abemaciclib, similar to what was observed in cells treated with the Mdm2 inhibitor nutlin-3a (Fig. 2C). Moreover, ChIP analyses revealed that abemaciclib enhanced p53 binding to the promoter regions of its target classical target gene *CDKN1A (p21)* and of the SASP factor *IGFBP3* (*26*), again similarly to what observed upon exposure to nutlin-3a (Fig. 2D). In accordance to the CDK4/6i-induced p53 activity and to the dependence of some SASP factors on this activity, abemaciclib (Fig. 2E) or palbociclib (Fig. 2F) treatment failed to induce expression of *LIF* and *IGFBP3* in BJ cells transduced with shRNA lentiviral particles against p53 (fig. S3, B and C). We then sought to validate the existence of a p53-dependent SASP in other senescence systems. First, we measured the levels of potential p53 SASP targets in a model of p53-induced but DDR-free senescence (*15*). BJ cells treated long-term with nutlin-3a showed high level of the classical p53 target genes *p21, GADD45* and *MDM2* (Fig. 2G) and an irreversible growth arrest (fig. S3D). Importantly, nutlin-3a-induced senescence was associated with a significant upregulation of the SASP factors *ISG15, IGFBP3, GDF15, TGFA* and *LIF* (Fig. 2G) while expression of NASP factors was not elevated (fig. S3E). To demonstrate the importance of p53 to promote part of the SASP in CDK4/6i-treated cells, we then made used of different systems and assays. Second, we measured the level of *LIF* and *IGFBP3* in genotoxic stress-induced senescent Mouse Dermal Fibroblasts (MDFs) with or without p53. Doxorubicin treatment failed to promote the transcription of *LIF* and *IGFBP3* in p53^-/-^ MDFs, but the induction was rescued in p53^-/-^ MDFs transduced with an expression vector carrying wild-type p53 (Fig. 2H). Together, these data show the existence of a SASP program positively regulated by the transcriptional activity of p53, from now on defined the p53-associated secretory phenotype (PASP).

**Figure 2.**
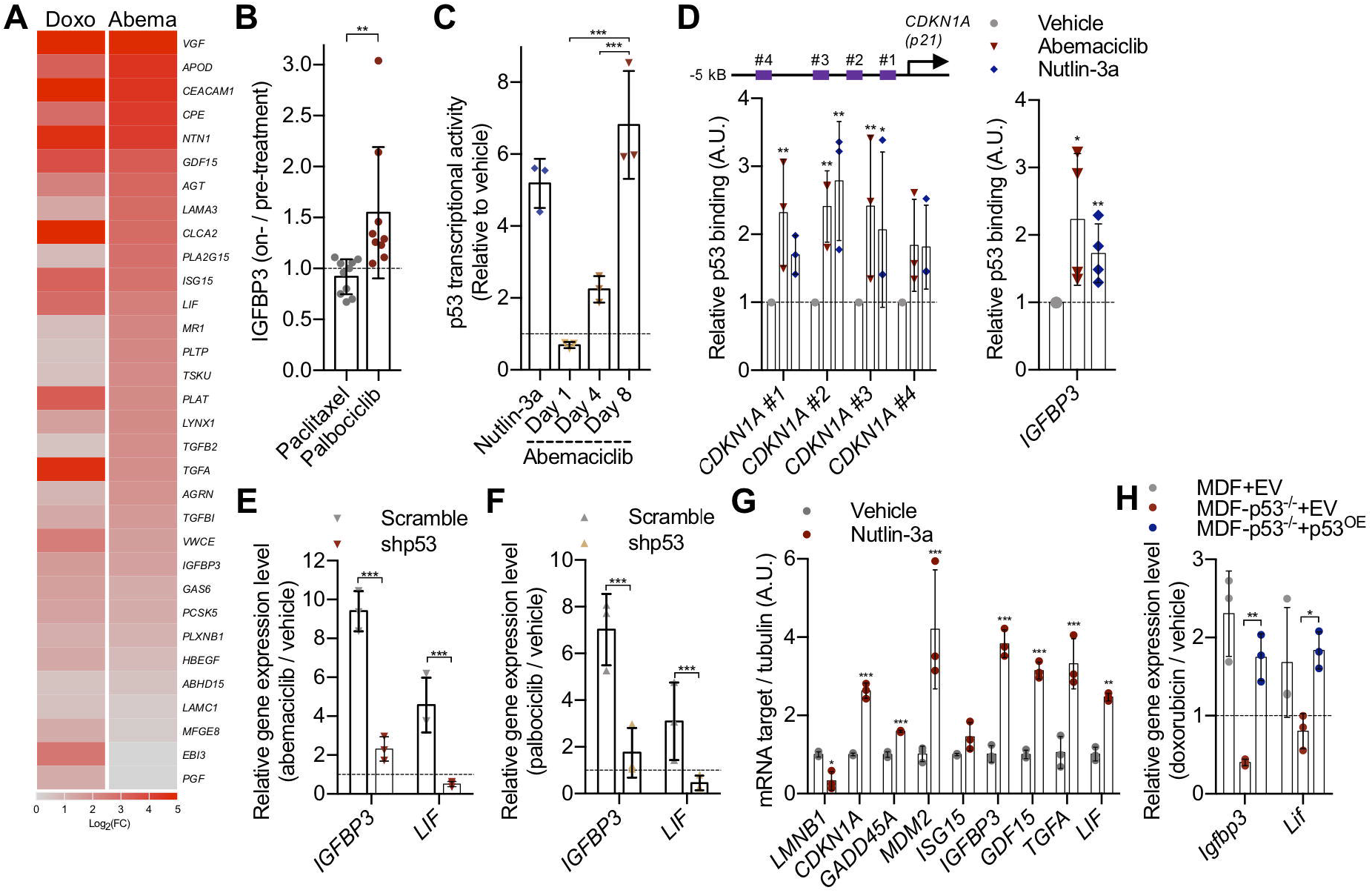
CDK4/6i treatment induces a p53-dependent SASP. (**A**) Human fibroblasts (BJ) were treated with vehicle (water for 8 times 24 hours), abemaciclib (1 μM for 8 times 24 hours) or doxorubicin (250 nM for 24 hours). RNA-sequencing was performed with treated cells at 8 dpt (N=3 independent samples, sequenced together) and heatmap of the p53-dependent SASP (PASP) for doxorubicin- or abemaciclib-treated groups (8 dpt) relative to vehicle-treated group from the RNA-sequencing datasets (p-value<0.01 was regarded as significant). (**B**) Cell-free plasma was derived from breast cancer patients treated with paclitaxel (N=10) or palbociclib (N=9) and the expression levels of IGFBP3 quantified by ELISA. The values represent the ratio between on-treatment and pre-treatment levels for each patient. (**C**) BJ fibroblasts transduced with a p53 reporter were treated with nutlin-3a (positive control) or abemaciclib (1 μM for 1 or 4 or 8 times 24 hours) and luciferase activity measured after treatments at the indicated time points (N=3). (**D**) Chromatin was extracted from BJ fibroblasts treated with vehicle or abemaciclib or nutlin-3a and ChIP assays using an antibody against p53 were performed. qRT-PCR was performed using primers amplifying the promoter region of *CDKN1A* or *IGFBP3* containing p53 binding sites. Values indicate fold enrichment relative to the vehicle group (N=4). (**E** and **F**) RNA was isolated from vehicle, abemaciclib (1 μM) (**E**) or palbociclib (1 μM) (**F**) treated scramble/shp53 BJ cells and quantified by qRT-PCR for *IGFBP3* or *LIF* genes (N=3). (**G**) RNA was isolated from BJ fibroblasts treated with vehicle (DMSO for 8 times 24 hours) or nutlin-3a (10 μM for 8 times 24 hours) at 8 dpt and quantified by qRT-PCR for *LMNB1* and p53 targets genes (N=3). (**H**) Mouse dermal fibroblasts (MDF) or MDF-p53^-/-^ fibroblasts were transfected with empty vector (EV) or p53-overexpressing vector (p53^OE^), then the cells were treated with vehicle (water) or doxorubicin (250 nM for 24 hours) and qRT-PCR was performed for *Igfbp3* and *Lif* (p53-dependent) genes (N=3). Unpaired two-tailed *t*-test, data are means ±SD (**B**). One-way ANOVA, data are means ±SD (**C** and **D**). Twoway ANOVA, data are means ±SD (**E**, **F**, **G** and **H**). *p<0.05, **p<0.01, ***p<0.001, N.S.=not significant. dpt, days post treatment.

### Senescent cells with a p53-dependent SASP only are well-tolerated *in vivo*

To evaluate the presence and tolerability of senescent cells with PASP but without NASP *in vivo,* we exposed cancer-free p16-3MR mice, which harbor a Renilla Luciferase (RL) reporter gene driven by the p16 promoter (*27*), to clinically-relevant doses of abemaciclib or doxorubicin (fig. S4, A to C). Abemaciclib- and doxorubicin-treated mice showed a similar enhanced bioluminescent signal (Fig. 3, A and B), and comparable induction of p16 and SA-β-gal in kidneys (Fig. 3, C to E). However, in alignment with cell culture measurements, doxorubicin treatment increased the concentration of CXCL1/KC in mouse plasma (Fig. 3F) and promoted expression of IL-6 (Fig. 3G) and various other NASP factors in kidney and liver (Fig. 3H and fig. S4D), while abemaciclib had no effects. In contrast, PASP factors *Isg15, Igfbp3, Gdf15, Tgfa* and *Lif* were equally upregulated in the kidneys of mice exposed to doxorubicin or abemaciclib (Fig. 3I). During treatment, and up to 14 days post, doxorubicin-treated mice showed substantial weight loss, while abemaciclib-treated mice did not lose any weight (Fig. 3J). Blood cell counts in the abemaciclib-treated group did not differ when compared to control animals, while doxorubicin caused anemia, as measured by reduced number of red blood cells (Fig. 3K), and an overall slight decrease in the total number of leukocytes (Fig. 3L) with impaired proportion of B cells, granulocytes and macrophages (Fig. 3M). As previously shown (*6*), treatment with doxorubicin severely affected strength and physical activity, as measured by rotarod assay (Fig. 3N), grip strength (Fig. 3O) and hanging endurance (Fig. 3P) at both 7 and 14 days after treatment. In contrast, mice treated with abemaciclib did not show any apparent reduction of physical performances in comparison to vehicle-treated cohorts (Fig. 3, N to P). Together, these data suggest that senescent cells with a PASP but without a NASP are generated *in vivo* but do not exert detrimental functions.

**Figure 3.**
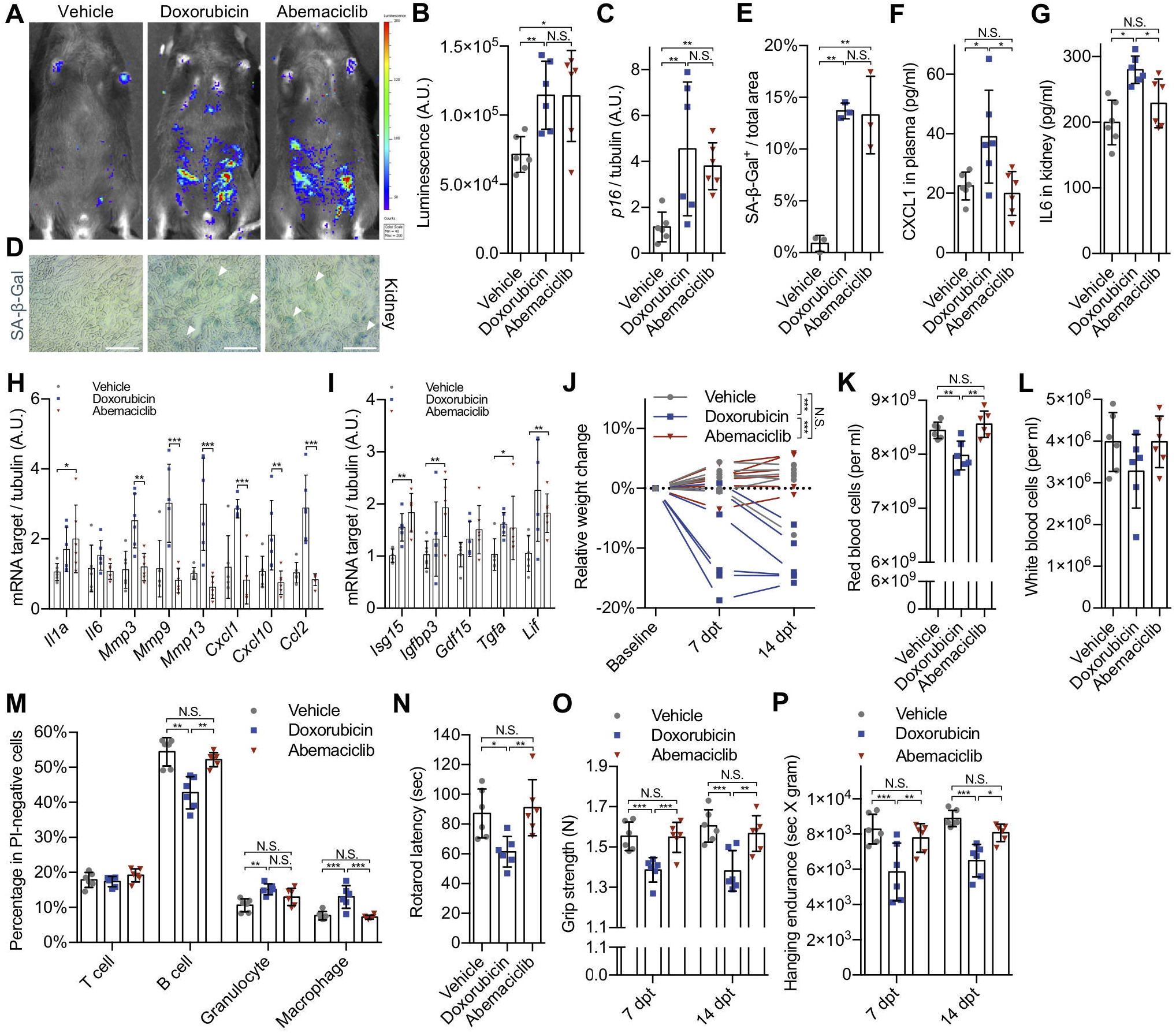
Senescent cells with a p53-dependent SASP only are well-tolerated *in vivo.* p16-3MR mice were treated with vehicle (PBS, 7 consecutive days), doxorubicin (5 mg/kg, 3 consecutive days) or abemaciclib (50 mg/kg, 7 consecutive days). N=6 mice/group. 14 dpt, bioluminescence was visualized and quantified by the IVIS spectrum *in vivo* imaging system, as shown by representative bioluminescence images (**A**) and quantification (**B**). (**C**) RNA isolated from treated kidneys and mRNA encoding p16 quantified by qRT-PCR. Representative images (**D**) to visualize SA-β-gal activities in vehicle-, doxorubicin-or abemaciclib-treated mouse kidney sections at 15 dpt (arrows indicated positive area; scale bar, 1 mm; N=3) and quantified (**E**). (**F**) 15 dpt, plasma was collected and expression levels of CXCL1 measured by ELISA. (**G**) Protein lysate obtained from drug treated kidneys to quantify IL6 by ELISA. (**H** and **I**) RNA isolated from kidneys and mRNA encoding indicated genes quantified by qRT-PCR. (**J**) Relative weight changes were calculated at 7 dpt and 14 dpt. Red blood cells (**K**) and white blood cells (**L**) were counted at 15 dpt. Percentage of T cells, B cells, granulocytes and macrophages were determined by flow cytometry analysis (**M**). Physical performance was measured by rotarods assay at 15 dpt (**N**), grip strength meter at 7 dpt and 14 dpt (**O**), and hanging tests were performed at 7 dpt and 14 dpt and normalized to weights (**P**). One-way ANOVA, data are means ±SD (**B, C, E, F, G, K, L** and **N**). Two-way ANOVA, data are means ±SD (**H, I, J, M, O** and **P**). *p<0.05, **p<0.01, ***p<0.001, N.S.=not significant. dpt, days post treatment.

### p53-associated SASP lacks pro-tumorigenic properties

Genotoxic stress-induced senescent cells become paradoxically pro-tumorigenic via secretion of various SASP factors (*5, 6*). To understand whether SASP factors part of the PASP can be pro-tumorigenic, we used different *ex vivo* and *in vivo* models. Exposure of MCF7 (breast cancer) (Fig. 4A), A549 and HCC827 (lung cancer) (fig. S5A) cells to the CM of abemaciclib-induced senescent cells had a strongly reduced pro-proliferative effect in comparison to CM from doxorubicin-induced senescent cells. Similarly, co-injection of doxorubicin-induced or abemaciclib-induced senescent fibroblasts (BJ) and lung cancer cells (A549) into the flanks of *Foxn1*^Nu^ mice indicated that doxorubicin-treated, but not abemaciclib-treated, cells promoted tumor growth (Fig. 4, B and C and fig. S5B). Moreover, CM from doxorubicin-treated BJ cells promoted migration of MCF7 and A549 cells while no significant effect was observed in cells exposed to CM from abemaciclib-treated cells (Fig. 4, D and E). To compare the effect on tumorigenesis of normal cells induced to senescence by CDK4/6i or genotoxic stress *in vivo*, we treated mice with abemaciclib and doxorubicin and 7 days later transplanted the syngeneic breast cancer cells MMTV-PyMT in the mammary fat pad. Mice that were treated pre-implantation with doxorubicin showed accelerated tumor growth (Fig. 4F) and reduced survival rate (Fig. 4G), while mice treated with abemaciclib did not show significant differences when compared to control animals. Our data show that senescent cells with PASP only do not acquire pro-tumorigenic properties.

**Figure 4.**
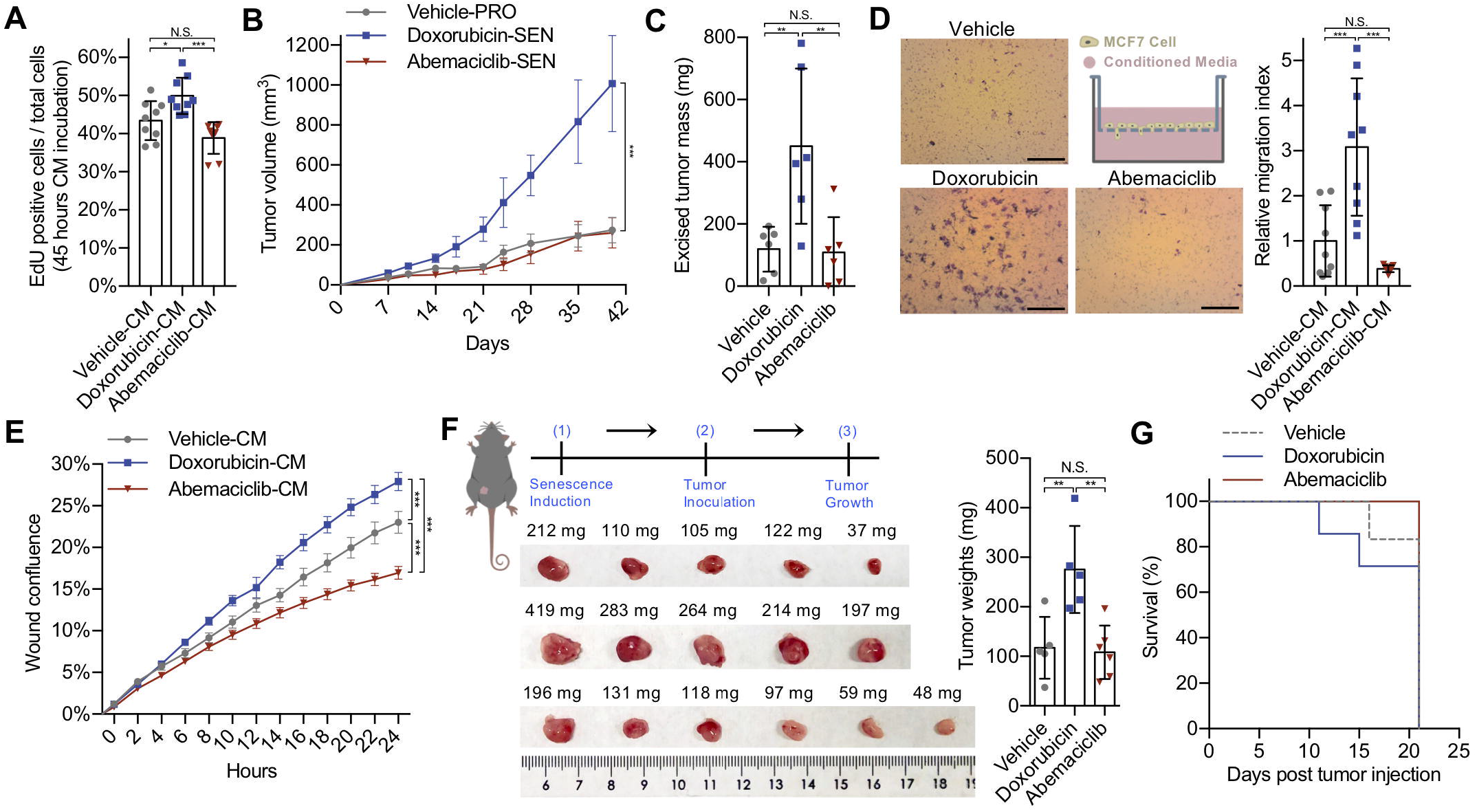
p53-associated SASP lacks pro-tumorigenic properties. (**A**) MCF7 cells were incubated with serum-free CM collected from treated BJ fibroblasts (8 dpt) containing EdU for 45 hours and EdU^+^ cells quantified (N=9). (**B** and **C**) Treated BJ fibroblasts (2.5×10^5^) were co-injected with A549 cancer cells (10^6^) subcutaneously in *Foxn1^Nu^* mice, tumor growth was measured at the indicated times (N=6 mice/group) (**B**) and the excised tumors were weighed (**C**). (**D**) MCF7 cells migrated through the pores of trans-well after 24 hours incubation with CM (8 dpt; scale bar, 1 mm; n=9) were stained with 0.2% crystal violet and quantified. (**E**) Migration of A549 cells exposed to CM collected from treated BJ cells (8 dpt; N=4) was evaluated using a scratch assay. Imaging and analysis were done using a live imaging system. (**F** and **G**) p16-3MR mice were treated with vehicle (PBS, 7 consecutive days), doxorubicin (5 mg/kg, 3 consecutive days) or abemaciclib (50 mg/kg, 7 consecutive days). N=6 or 7 mice/group. At 14 dpt, MMTV-PyMT mouse breast cancer cells were implanted in the mammary fat pad of treated female mice. The tumors were excised and weighed 21 days post inoculation (left panel, images of the tumors; right panel, tumor weights) (**F**). The survival curves for different groups was plotted (**G**). One-way ANOVA, data are means ±SD (**A, C, D** and **F**). Two-way ANOVA, data are means ±SD (**B** and **E**). *p<0.05, **p<0.01, ***p<0.001, N.S.=not significant. dpt, days post treatment. PRO, proliferating cells. SEN, senescent cells. CM, conditioned media.

### Senescent cells with PASP and without NASP induces paracrine senescence and are effectively cleared

Cellular senescence can exert tumor suppressive non-cell autonomous functions by inducing paracrine senescence. To evaluate whether senescent cells with a PASP but without a NASP maintain paracrine senescence abilities, we injected wild-type senescent MDFs induced to senescence by doxorubicin or abemaciclib into the dorsal skin of p16-3MR mice. Interestingly, both subsets of senescent cells were equally able to promote luminescence in the transplanted area of the recipient mice, suggesting induction of p16^+^ cells (Fig. 5, A and B). Importantly, luminescence was not promoted by injection of non-senescent MDFs, confirming the validity of this *in vivo* assay for measuring paracrine senescence (Fig. 5, A and B). One of the major drivers of detrimental functions of senescent cells is their impaired turnover due to higher induction rates and lower immune-mediated clearance. No significant differences in the induction and accumulation of senescence were observed between the abemaciclib and doxorubicin groups at the time points chosen for phenotypical analyses (Fig. 3, A to E). To better characterize clearance rates, we decided to use a transplantation system that could compare removal of different subsets of senescent cells in the same immunological environment. MDFs derived from the p16-3MR mouse were induced to senescence *ex vivo* with doxorubicin or abemaciclib, and then transplanted into opposite flanks of immunocompetent and syngeneic wild-type untreated mice (Fig. 5C). Unexpectedly, absence of the pro-inflammatory NASP did not delay clearance, and abemaciclib-induced senescent cells showed even a faster removal compared to the doxorubicin-treated cells (Fig. 5C). Altogether, these data suggest that senescent cells with a PASP but without pro-inflammatory NASP retain the ability to promote paracrine senescence and are removed from tissues with faster kinetics.

**Figure 5.**
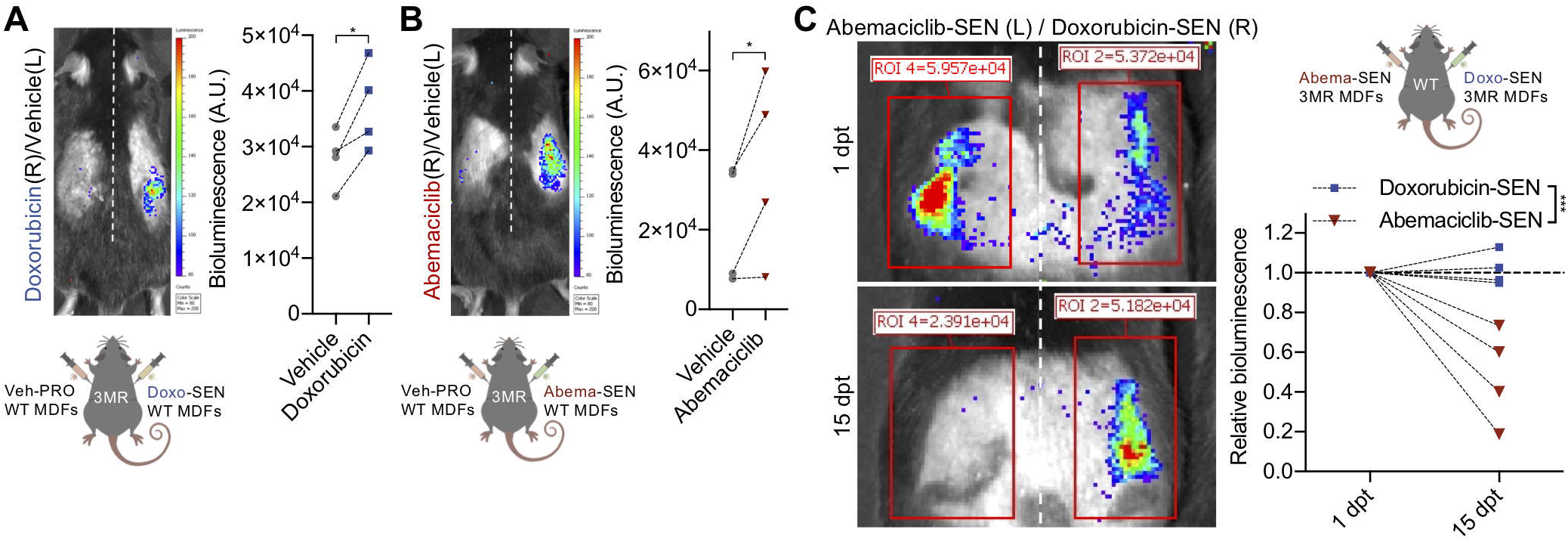
Senescent cells with PASP and without NASP induces paracrine senescence and are effectively cleared. (**A** and **B**) 2*10^5^ cells of vehicle-treated (water for 8 times 24 hours) mouse dermal fibroblasts (MDF) were injected into the left flank of the p16-3MR mice. Doxorubicin-induced (250 nM for 24 hours) (**A**) or abemaciclib-induced (4 μM for 8 times 24 hours) (**B**) MDFs were injected into the right flank of the same animal. 7 dpt, the above-mentioned mice were injected with coelenterazine and bioluminescence from the p16-3MR mouse was visualized/quantified by the IVIS spectrum *in vivo* imaging system and quantified. Lower panel, scheme of experimental design. (**C**) Equal amount (2*10^5^ cells) of doxorubicin-induced (250 nM for 24 hours, right flank) and abemaciclib-induced (4 μM for 8 times 24 hours, left flank) senescent MDF-3MR cells were subcutaneously injected into the same wild-type mice. At 1 dpt and 15 dpt, the above-mentioned mice were injected with coelenterazine and bioluminescence from the injected MDF-3MR cells was visualized/quantified by the IVIS spectrum in vivo imaging system and quantified. Upper panel, scheme of experimental design. Paired student *t*-test. Error bars, ±SD. *p<0.05, ***p<0.001. PRO, proliferating cells. SEN, senescent cells. dpt, days post treatment (transplantation).

## Discussion

Collectively, we show here show that CDK4/6i-treated non-malignant cells enter a senescent program characterized by morphological changes, suppression of many cell cycle-related genes, induction of a core senescence-associated signature and activation of lysosomal content and enzymes, but lacking many of the common SASP factors. This observation is in agreement with previous reports showing that overexpression of the CDK4/6i p16 leads to a senescent state with lower SASP (*13*). However, recent reports have highlighted that differences in SASP expression might not be only quantitative but also qualitative (*28, 29*).

Indeed, SASP composition might be dependent on various factors, including tissue, cell type and time point, and various signaling pathways including NF-κB, mTOR, cEBPβ, p38 and p53 (*30–32*). p53 signaling differs from the others because it was shown to interfere with other SASP programs, in particular via inhibition of NF-κB (*15*) and mTOR (*33*) signaling pathways. However, during the characterization of the CDK4/6i-induced senescence, we were able to identify a SASP transcriptional program that is promoted by p53 activity (PASP). Treatment with CDK4/6i induces a PASP and not a NASP, which reflects the lack of chronic DDR and activation of NF-κB signaling, two essential regulators of the pro-inflammatory SASP (*11, 12*). Prolonged abemaciclib treatment causes premature senescence *in vivo,* but the abemaciclib-induced senescent cells were well tolerated suggesting that the PASP does not exert detrimental functions. These data fit with the notion that toxicity of CDK4/6i in patients is generally lower than upon treatment with standard chemotherapy (*34*), and suggest that CDK4/6i could be better tolerated by cancer populations that are at high risk of developing side effects. Moreover, the presence *in vivo* of p16^+^ senescent cells with no SASP and detrimental effects emphasize how heterogeneous the senescence phenotype can be, and the importance of carefully phenotyping senescent cells to determine their potential toxicities. According to our data, one of the regulators of senescence-associated toxicities is predicted to be the NASP, while senescent cells with only PASP might be tolerated and potentially less dangerous. Thus, we suggest that a subset of PASP and NASP factors could be used as biomarkers for the potential detrimental function of senescent cells *in vivo*, and also as readouts for senotherapies. Because a major driver of deleterious effects of senescent cells is their aberrant persistence in tissues due to impaired removal, it is interesting to observe that cells with PASP-only are cleared at faster rates than cells with PASP and NASP. However, the SASP have variable and complex immune modulating effects (*35–38*) and it remains to be assessed whether the NASP actively interferes with immune-mediated clearance or if additional factors are involved. We also show that cells with PASP can transmit paracrine senescence, which could be a potent non-cell autonomous mechanism to extend tumor suppression. It remains to be elucidated if PASP-associated induction of paracrine senescence can have detrimental consequences in contexts other than cancer therapy, for example in aged tissues.

## Materials and Methods

### Cell culture and drug administration

BJ (CRL-2522), WI38 (CRL-7728), MMTV-PyMT (CRL-3278), A549 (CCL-185), HCC827 (CRL-2868), MCF7 (HTB-22) and hTERT-RPE1 (CRL-4000) cells were purchased from ATCC. Human MSCs were a gift from Prof. Irene Heijink (University Medical Center Groningen, The Netherlands). MEFs were generated from 13.5-day wild-type and p16^-/-^ embryos. MDFs were isolated from the dorsal skin of 3-month-old p53-null mice or wild-type littermates and a gift from Prof. Paul Hasty (University of Texas Health Science Center at San Antonio, USA). Cells were not re-authenticated by the laboratory but were regularly monitored for mycoplasma contaminations (once per month). No cell line used was listed in the database of commonly misidentified cell lines maintained by ICLAC. All cells were cultured in DMEM-GlutaMAX (Thermo Fisher) medium supplemented with 10% fetal bovine serum (GE Healthcare Life Sciences) and 1% penicillin-streptomycin (Lonza). All the human and mouse normal primary cells were maintained in 5% O_2_, 5% CO_2_, 37 °C incubators and all cancer cells were maintained in 20% O_2_, 5% CO_2_, 37 °C incubators. For drug treatment experiments, cells were plated in 6-well dishes with 3 technical repeats (3*10^4^ cells/well, around 30% confluence). Doxorubicin hydrochloride (Tebu-bio, BIA-D1202-1) was dissolved in sterile milliQ water at 250 μM as stock. Paclitaxel (MedChemExpress, HY-B0015) was dissolved in DMSO at 10 mM as stock. Palbociclib isethionate (Sigma-Aldrich, PZ0199) was dissolved in sterile milliQ water at 50 mM as stock. Abemaciclib (MedChemExpress, HY-16297) was dissolved in sterile milliQ water at 50 mM as stock. Nutlin-3a (MedChemExpress, HY-10029) was dissolved in DMSO at 10 mM as stock. All the drugs were further diluted in above-mentioned DMEM to treat cells at different concentrations as indicated in each figure legend.

### Colony formation assay

Drug-treated normal or cancer cells were re-plated in a 6-well (3*10^3^ cells/well) at the end of treatment, and allowed to grow in drug-free normal medium for 8 days. Cancer cells were plated in a 6-well (3*10^4^ cells/well), and incubated with serum-free conditioned media (CM) for 45 hours. Afterwards, cells were fixed in 4% PFA for 30 minutes and then stained with 0.2% crystal violet in 37% methanol for 1 hour. Pictures of all the plates were taken with a scanner (Epson). The images were cropped and processed in Microsoft PowerPoint using the same settings.

### EdU staining

Drug-treated cells were re-plated on coverslip in a 24-well plate (3*10^4^ cells/well) and cultured for 10 or 20 hours in the presence of EdU (10 μM), then fixed and stained as previously described (*39*). Images were acquired at 100 times magnification (Leica), and the number of cells was counted with the software ImageJ. The images were cropped using Microsoft PowerPoint.

### Senescence-associated β-galactosidase staining

For staining of cells, drug-treated cells were re-plated in a 24-well plate (2×10^4^ cells/well), and the SA-β-galactosidase staining was done as described previously (Hernandez-Segura et al., 2018a). For *in vivo* kidney section staining, kidneys from drug-treated and control animals were immediately imbedded in O.C.T (Sakura, 621232), snap-frozen in liquid nitrogen and stored in −80 °C. The sectioning (10 μm) was done using cryostat one hour prior to staining, the slides washed with prechilled PBS on ice in a glass coplin jar twice for 5 minutes and fixed in a mixture of formaldehyde (2%) and gluteraldehyde (0.2%) for 10 minutes on ice. After fixation, the slides were washed briefly with milliQ water, and then stained with the staining buffer overnight in 37 °C incubator without CO_2_. The staining buffer included 1 mg/ml X-gal (Thermo Fisher) in dimethylformamide, 40 mM citric acid/Na phosphate buffer, 5 mM potassium ferrocyanide, 5 mM potassium ferricyanide, 150 mM sodium chloride and 2 mM magnesium chloride. Next day, the staining solution was washed away with milliQ water and the slides were mounted with 70% glycerol. Images were acquired at 100x magnification with a microscope (Leica) and were processed in Microsoft PowerPoint. The SA-β-gal positive area and total area was measured and quantified by with the software ImageJ.

### Immunofluorescence

Cells were re-plated in 24-well dishes with coverslip (3×10^4^ cells/well) and cultured in normal medium for 8 days. On the day of staining, the cells were gently washed with PBS for 3 times, fixed 10-15 minutes with 4% PFA, washed 3 times with PBS and permeabilized for 15 minutes with 0.1% Triton-100-X in PBS. After 3 washes with PBS, cells were blocked for 45 min in 3% BSA/PBS and incubated in primary antibody (gamma H2AX, 1:800 dilution; Novus, NB100-384) in 3% BSA/PBS at 4 °C overnight. After 3 washes with PBS, coverslips were incubated with secondary antibody for 45 minutes in dark at room temperature and then with 2 μg/ml DAPI for 15 minutes in dark at room temperature. After washing 3 times, the samples were mounted with ProLong Gold Antifade Mountant (Thermo Fisher). Images were acquired at 100X magnification with a microscope (Leica).

### Conditioned media (CM) collection and analyses

BJ fibroblasts were treated with vehicle (water − 8 times 24 hours; 1 in 1,000), doxorubicin (250 nM – 1 time 24 hours) or abemaciclib (1 μM – 8 times 24 hours). After the treatments, cells were cultured with drug-free normal medium for 8 days. Thereafter, cells from different groups were trypsinized and counted, and 10^6^ cells from each group were re-plated and incubated with serum-free medium for 24 hours. Then the CM was collected and centrifuged at 300 g / 5 minutes to remove any floating cell or debris. The CM was kept in −80 °C after harvesting and thawed on ice for further analyses or experiments. Human Cytokine Antibody Array (Abcam, ab133997) was used following manufacturer’s instructions. After the images were taken, density of positive controls from 4 membranes was equalized using Adobe Photoshop, and individual factors from different membranes compared. For ELISA, human CXCL1/GROα duo-set or CCL5 duo-set or IGFBP3 duo-set (R&D Systems) were used to detect the concentrations of CXCL1 or CCL5 or IGFBP3 in the CM following manufacturer’s instructions.

### Western blot

Total cell lysates were obtained by resuspension of the cells in RIPA buffer (cat# ab 156034, Abcam) supplemented with proteinase and phosphatase inhibitors (cat# A32959, Pierce (TFS)). Protein concentration was measured using a BCA protein assay kit (cat# 10741395, Pierce (TFS)) and subsequently protein sample buffer was added to a final concentration of 40 mM Tris-HCl (pH 6.8), 1.9% SDS, 3.1% β-mercaptoethanol and 6,3% glycerol. Samples were boiled for 5 minutes. Equal amount of proteins was loaded onto 12% SDS-PAGE gel (acrylamide: bisacrylamide 29:1, cat# 161-0146, Bio-Rad) and after size separation blotted onto 0.2 μm nitrocellulose membrane (cat# 162-0112, Bio-Rad). Immunodetection was performed by standard procedures of p16/Ink4a (clone EPR1473, cat# ab108349, Abcam), p21 (clone C-19, cat# sc-397, Santa Cruz Biotechnology), p53 (clone DO-1, cat# sc-126, Santa Cruz Biotechnology), Rb (clone 4H1, cat# 9309, Cell Signaling Technology) and phospho-Rb (ser795, cat# 9301, Cell Signaling Technology). The antibodies for the proteins of interest were used in 1:1000 dilution in 5%milk/TBST (Tris-buffered Saline with 0.1% Tween20) and incubated overnight at 4 degrees Celsius. Immunodetection of vinculin (cat# V9131, Sigma-Aldrich) or β-Actin (clone C4, cat# 08691001, MP Biomedicals) was performed as loading control. ECL Prime western blotting detection reagent (cat# RPN2232, GE Healthcare) was used according to manufacturer’s guidelines for detection and the signal was measured using an ImageQuant LAS 4000 biomolecular imager (GE Healthcare). Densitometry was performed using ImageJ software (NIH – Public domain) and values were corrected for protein input as measured by the loading control.

### Generation of a lentiviral NF-κB reporter vector

Plasmid pGL4.32[luc2P/NF-κB-RE/Hygro] (cat# E8491, Promega) was used as template DNA in a PCR for obtaining the minimal promoter Luc2P together with the NF-κB response elements (NF-κB-RE). The DNA polymerase used was Phusion High Fidelity (cat# E0553S, New England Biolabs (NEB)). Forward primer: 5’-aaaaATCGATgg cctaactggccggtacc-3’, reverse primer 5’-aaaaGGATCCcgactctagagtcgcggcc-3’. The PCR product was purified from gel cut with ClaI (cat#10656291001, Roche) and BamHI-HF (cat# R3136s, NEB) overnight at 16 degrees Celsius. This fragment was ligated into the backbone vector pLenti CMV GFP Hygro (656-4) (restriction with ClaI and BamHI-HF released the CMV promoter from the lentiviral backbone sequence). pLenti [CM V/GFP/Hygro] (656-4) was a gift from Eric Campeau & Paul Kaufman (Addgene plasmid #17446). The cut backbone vector was treated with Antarctic Phosphatase (cat# M0289S, NEB) before ligation. T4 DNA ligase (cat# M0202S, NEB) was used in the ligation reaction. All enzymes were used according to manufacturer’s manual. The ligation reaction products were transformed into NEB stable Competent *E. coli* (High Efficiency, cat# c3040h, NEB) and grown overnight at 37 degrees Celsius on LB plates supplemented with 50 μg/ml Ampicillin. A colony PCR was performed to check for positive clones. Single positive clones were grown overnight in liquid LB supplemented with Ampicillin at 37 degrees Celsius to be midiprepped the next day using a PureLink^®^ HiPure Plasmid Midiprep Kit (cat# K210005, Invitrogen). This newly generated lentiviral NF-κB-reporter vector (pLenti-NF-κB-RE-minP-LUC-EGFP-hygro) was send to GATC/Eurofins for sequencing and verification of the Luc2P/ NF-κB-RE insert. Recombinant human TNF-α (PeproTech, 300-01A) (10 ng/μl; 6 hours treatment) was used as positive control to validate the NF-κB reporter.

### Generation of a lentiviral p53-reporter vector

Plasmid PG13-luc (wt p53 binding sites) was used as template DNA in a PCR for obtaining the p53 wild-type binding sites together with the luciferase reporter. PG13-luc (wt p53 binding sites) was a gift from Bert Vogelstein (Addgene plasmid # 16442). The DNA polymerase used was Phusion High Fidelity (cat# E0553S, NEB). Forward primer 5’-aaaaaGGGACCCaaaCGATAAGCTTGATGCC-3’, reverse primer 5’-tttttGTCGACaaaTTAAATCTC TGTAGG-3’. The PCR product was purified from gel and cut with KflI (cat# FD2164, ThermoFisher Scientific) and SalI (cat# R3138, NEB) for 15 minutes at 37 degrees Celsius. This cut PCR product was ligated into the backbone vector pLenti CMV GFP Hygro (656-4) (restriction with KflI ans SalI releases the cPPT/CST element, the CMV promoter and the EGFP from the lentiviral backbone sequence. The cut backbone vector was treated with Antarctic Phosphatase (cat# M0289S, NEB) before ligation. T4 DNA ligase (cat# M0202S, NEB) was used in the ligation reaction. All enzymes were used according to manufacturer’s manual. The ligation reaction products were transformed into NEB stable Competent *E. coli* (High Efficiency, cat# c3040h, NEB) and grown overnight at 37 degrees Celsius on LB plates supplemented with 50 μg/ml Ampicillin. A colony PCR was performed to check for positive clones. Single positive clones were grown overnight in liquid LB supplemented with Ampicillin at 37 degrees Celsius to be midiprepped the next day using a PureLink^®^ HiPure Plasmid Midiprep Kit (cat# K210005, Invitrogen). This newly generated intermediate lentiviral p53-reporter vector (pLenti–p53 WT bind. sites-LUC-hygro) was still dysfunctional because of lacking the cPPT/CST element. The cPPT/CST element has been repaired before the newly generated lentiviral p53-reporter vector (pLenti–p53 WT bind.sites-LUC-hygro) was sent to GATC/Eurofins for sequencing and verification of the cPPT/CST-p53 WT bind.sites-LUC insert. Nutlin-3a (MedChemExpress, HY-10029) (10 μM; 24 hours treatment) was used as positive control to validate the p53-reporter.

### Production of lentivirus

293FT cells were seeded in a 10 cm petridish at 6*10^6^ cells. Next day, cells were transfected with the ViraPower plasmid mix (cat# K4970-00, Invitrogen) together with the lentiviral vectors using PolyFect transfection reagent (cat# 301105, Qiagen) overnight. The medium with transfection mix was replaced for normal growth medium without antibiotics, collected after 24 hours and concentrated using Peg-It Virus precipitation solution (cat# LV810A-1, System Biosciences) according to the manufacturer’s manual.

### Lentivirus transduction

Lentiviral particles were either generated by the laboratory (NF-κB or p53 reporter) or obtained from the MISSION shRNA library (Sigma-Aldrich). These obtained lentiviral particles harbor the following shRNA clones in pLKO.1 backbone vector and were used to infect BJ cells: p53 (*TP53*)–TRCN0000003755. Titers of these lentiviral particles were all equal to or higher than 1.8E+07 TU/ml. 10^5^ BJ cells per well were seeded in a 6 well plate format on day 0. On day 1, cells were infected with the lentiviral particles (1:20 dilution in complete medium) for 24 hrs. Polybrene (6 μg/ml; cat# sc-134220, Santa Cruz Biotechnology) was added together with the lentiviral particles to enhance the infection efficiency. Cells were washed and refreshed with complete medium on day 2. Selection with puromycin (2 μg/ml; cat# P8833, Sigma-Aldrich) started on day 3 for 3 days. BJ cells infected with pLKO.1-scrambled lentiviral particles were used to serve as control cells.

### RNA sequencing assay

Total RNA was extracted from treated cells via an ISOLATE II RNA mini kit (Bioline, Cat: BIO-52073) following manufacturer’s instructions. The extracted RNA was quantitated using a Nanodrop and RNA quality was measured via BioAnalyzer RNA chip (Agilent). Poly-A tail selection was used to enrich for messenger RNA (mRNA) using the NEXTflex Poly(A) Beads kit (Bioo Scientific Corp, cat #512980). RNA-Seq library preparation was carried out using NEXTflex Rapid Directional qRNA-Seq kit (Bioo Scientific, cat# 5130-05D). In brief, mRNA was fragmented using a cationic buffer and then submitted to first and second strand synthesis, followed by adenylation, ligation and 15 cycles of PCR. Library quality and size distribution were validated on a Qubit (Thermo Fisher Scientific) and an Agilent High Sensitivity DNA chip. Clusters for sequencing were generated on the cBot (Illumina). Paired-end sequencing was performed at 400 M reads per lane of the flow cell on an Illumina HiSeq 2500. Average quality scores for the completed run across all samples was >30, with an average of 10 million reads for each pooled sample. The read length was 76 bp. Raw sequencing data were demultiplexed according to library-specific barcodes and converted to fastq format using standard Illumina software (bcl2fastq version 1.8.4). The resulting reads were mapped to the human reference genome (GRCh38) using Bowtie2 (version 2.2.4). Sequencing data is available on ArrayExpress under accession no. **E-MTAB-7642**. For differential gene expression, we used DESeq2 (*40*) to evaluate the genes that were differentially expressed among all the different treatments compared to the vehicle-treated control. We considered those genes with an adjusted p-value lower than 0.01 as being differentially expressed. GraphPad Prism 7 was used to generate the heat-map using the logarithm base 2 of fold change. Enriched pathways in the differentially expressed genes were evaluated using the online tool “Over-representation analysis” of the Consensus Path DB-human (http://cpdb.molgen.mpg.de/) (*41*). For the Venn plots showing similarly expressed genes in different conditions, only genes that were differentially expressed with a fold change in the same direction (up-or down-regulated in samples compared) were considered. Venn plots were generated in R version 3.5.1 using the packages “VennDiagram”.

### qRT-PCR assay

Total RNA was isolated using an Isolate II Rna Mini Kit (Bioline). 250~500 ng of RNA was reverse transcribed into cDNA using the High-Capacity cDNA reverse transcription kit (Applied Biosystems). qRT-PCR reactions were performed using the Universal Probe Library system (Roche) and a SensiFast Probe kit (Bioline). Expression of tubulin was used to normalize the expression of target genes. Every biological replicate was analyzed in duplicate. List of primers is provided in **supplementary table S2**.

### Chromatin immunoprecipitation (ChIP) assay

BJ cells treated with vehicle, abemaciclib (1 μM – 8 times 24 hours) or nutlin-3a (10 μM – 8 times 24 hours) were crosslinked and processed using the ChIP kit (ab500, Abcam) following manufacturer’s instructions. 6 rounds of sonication (high energy, 30 seconds on / 30 seconds off) was done using a sonicator. After DNA was purified, qRT-PCR was done with SYBR green super mix (1725120, Biorad) using primers as followed:

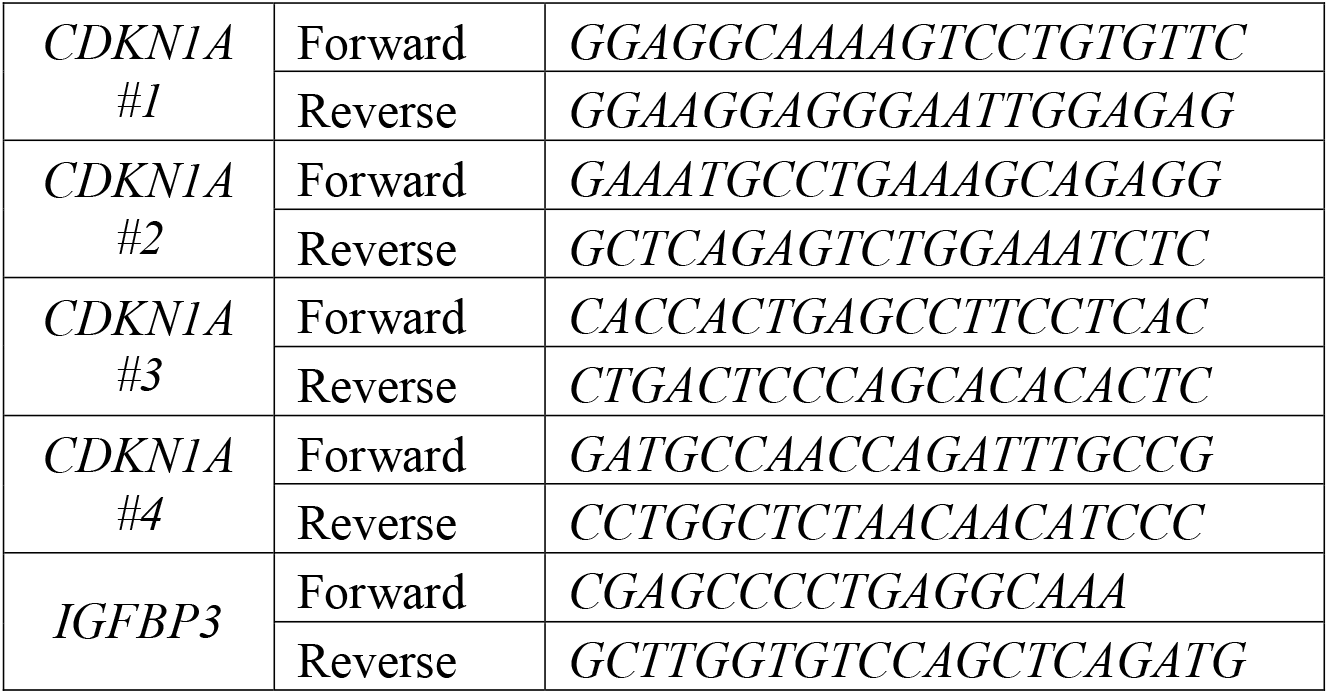

### Transwell migration assay

MCF7 breast cancer cells (2*10^4^) were re-suspended in serum-free medium and seeded in a transwell insert (8 μm pore, Corning) with serum-free media collected from drug-induced senescent fibroblasts in the outer chamber of a 24-well culture dish. Cells were cultured in a 5% O_2_, 5% CO_2_, 37 °C incubator for 16 hours. Thereafter, cells migrated to the lower side of the trans-well were fixed with 4% PFA and stained with 0.2% crystal violet. Images were acquired at 40X magnification with a microscope (Leica).

### Scratch migration assay and IncuCyte imaging

A549 lung cancer cells (4*10^4^) were plated in a 96-well ImageLock plate (Sartorius) and cultured in a 20% O_2_, 5% CO_2_, 37 °C incubator for 24 hours. Then the 96-pin IncuCyte WoundMaker was used to create precise and reproducible cell-free zone in each well. Cells were washed with PBS and then incubated with serum-free conditioned media collected from drug-induced senescent fibroblasts for 24 hours in the IncuCyte imaging machine. Images were taken every 2 hours and analyzed by the IncuCyte software.

### *In vivo* animal experiments

All the mice were maintained in the central animal facility (CDP) of University Medical Center Groningen (UMCG) under standard conditions. All the experiments were approved by the Central Authority for Scientific Procedures on Animals (CCD – License #AVD105002015339 and #AVD1050020184807) in the Netherlands.

For abemaciclib anti-tumor effect validation, 10^6^ fLUC-MMTV-PyMT cancer cells, previously generated (Demaria et al., 2017), were injected into the mammary fat pad of 13-week-old female p16-3MR mice under anesthesia (10 mice). At 7 days and 14 days post cell injection, the mice were injected with 150 mg/kg D-Luciferin (Sanbio, 14681-1). 8 minutes later, the mice were anesthetized with 2% isoflurane and firefly bioluminescence was visualized/quantified using the IVIS Spectrum *In Vivo* Imaging System (PerkinElmer, 5-minute medium binning) in CDP-UMCG. The mice were equally distributed in two groups based on tumor size (bioluminescence).

The female mice bearing breast cancer were then injected i.p. with vehicle (PBS, 7 consecutive days) or abemaciclib (in PBS, 50 mg/kg, 7 consecutive days) (N=5 mice/group). After the second bioluminescence imaging, the mice were terminated, and tumors were excised for weighing.

For senescence induction and healthspan analysis in p16-3MR, both male and female mice were used. 14-week-old healthy mice were injected i.p. with vehicle (PBS, 7 consecutive days), doxorubicin (in PBS, 5 mg/kg, 3 consecutive days) or abemaciclib (in PBS, 50 mg/kg, 7 consecutive days) (N=6 mice/group). All the treatments were finished at the same day. At 14 days post above-mentioned treatments, the mice were injected with 100 μl Xenolight RediJect Coelenterazine h (PerkinElmer, 760506). 20 minutes post injection, mice were anesthetized with 2% isoflurane and renilla bioluminescence was visualized/quantified by the IVIS Spectrum *In Vivo* Imaging System (PerkinElmer, 5-minute medium binning) in CDP-UMCG. Forelimb grip strength (N) was measured using a grip strength meter (Columbus Instruments) at 7 and 14 days post treatments, and results were averaged from 5 trials. Hanging tests were performed at 7 and 14 days post treatments. The mice were placed under the grid of above-mentioned grip strength meter with the bedding below to protect the mice when they fell down. A cut-off time of 300 seconds was determined, and the hanging time was normalized to body weights as hanging duration (sec) multiplied by body weight (g). At 15 days post treatments, endurance tests were performed on an accelerating RotaRod (IITC Life Science) using a top speed of 40 r.p.m. over a period of 300 seconds. Each mouse was put on the machine to adapt twice, four hours later two trials were recorded for each mouse. The average of the two trials was calculated and plotted. For *ex vivo* analysis, kidneys and livers were both snap-frozen in liquid nitrogen for RNA/protein isolation and imbedded in O.C.T (Sakura, 621232) (frozen in liquid nitrogen) for cryo-sectioning. All the samples were stored in −80 °C until analyzed. 20 μg total protein was collected from each treated kidney and IL6 protein level was determined by the mouse IL6 duo-set (R&D Systems) ELISA. The experiments were done following manufacturer’s instructions. Part of the plasma was used for *in vivo* ELISA, 100 μl plasma from each mouse was tested using the mouse CXCL1/KC duo-set (R&D Systems). The experiments were done following manufacturer’s instructions. Around 25 μl of plasma was diluted and used for red blood cell and white blood cell counting (Medonic CA620). Around 900 μl of plasma from each mouse was used to stain for T cell (1:150, APC-CD3e/clone 145-2C11, Biolegend Cat. 100312), B cell (1:300, FITC-CD45R/B220/clone RA3-6B2, Biolegend Cat. 103205), granulocyte and macrophage (1:1000, PECy7-Gr-1/clone RB6-8C5 and PECy7-CD11b /clone M1/70, Biolegend Cat.108416 and Cat.101216), then analyzed by flow cytometry (BD MoFlo XDP).

For MMTV-PyMT cancer cell injection in doxorubicin/abemaciclib-treated mice, 14-week-old healthy female mice were injected i.p. with vehicle (PBS, 7 consecutive days; N=6 mice), doxorubicin (in PBS, 5 mg/kg, 3 consecutive days; N=7 mice) or abemaciclib (in PBS, 50 mg/kg, 7 consecutive days; N=6 mice). All the treatments were finished at the same day. At 7 days post above-mentioned treatments, 3*10^5^ fLUC-MMTV-PyMT cancer cells were injected into the mammary fat pad of the treated female mice under anesthesia. The tumors were allowed to grow for 21 days. Then the mice were terminated, and tumors were excised for weighing and photos.

For human senescent/cancer cells co-injection in *Foxn1*^Nu^ mice, BJ fibroblasts were treated with vehicle (water for 8 times 24 hours; 1 in 1,000), doxorubicin (250 nM for 24 hours), palbociclib or abemaciclib (both 1 μM for 8 times 24 hours) *in vitro.* At 8 days post drug removal, BJ cells (2.5*10^5^) from each group were co-injected with A549 cancer cells (10^6^) subcutaneously in right flanks of 12-week-old female *Foxn1*^Nu^ mice (The Jackson Laboratory)(N=6 mice/group). 7 days after cell injection, tumor volume was calculated twice a week. The tumor length, width and depth were measured with a caliper and the investigator was blinded. The mice were terminated when one of the tumors reached the size for humane endpoint (>1,500 mm^3^). After the tumors were excised, the weight of each tumor was measured and plotted.

### Patient characteristics and plasma analysis

Paired plasma samples before start of therapy (pre-treatment) and after ±4 weeks (median 29 days) of treatment (on-treatment) were obtained from 9 metastatic breast cancer patients who received palbociclib treatment and 10 metastatic breast cancer patients who received paclitaxel treatment. Written informed consent was obtained from all patients. Mean age was 55.9 years (SD 10.9) for the palbociclib group and 60.1 years (SD 7.4) for the paclitaxel group. The samples were collected in cell stabilizing blood collection tubes (CellSave or Streck Cell-Free DNA BCT tubes). After collection the blood was centrifuged twice (1700 g for 10 minutes and 12000 g for 10 minutes). Afterwards the plasma was pooled and directly stored in 2 ml tubes in −80 °C until analyzed. For ELISA, human CXCL1/GROα duo-set or CCL5 duo-set or IGFBP3 duo-set (R&D Systems) were used to detect the concentrations of CXCL1 or CCL5 or IGFBP3 in the plasma following manufacturer’s instructions.

### Statistics

Graphpad Prism 7 was used for the statistical analyses. Detailed information is provided in each figure legend.

## Acknowledgments

We thank Bertien Dethmers and Gerald de Haan for help with blood analysis. We are thankful to the Demaria laboratory for fruitful discussions.

## Funding

This work was supported by the University Medical Center Groningen and by a grant from the Dutch Cancer Foundation (KWF) to M.D.

## Notes

### Competing Interest Statement

M.D. is founder and shareholder of Cleara Biotech

