## Supplementary Figures 1-6 for "Pharmacological CDK4/6 inhibition unravels a p53-induced secretory phenotype in senescent cells"

**Title:**

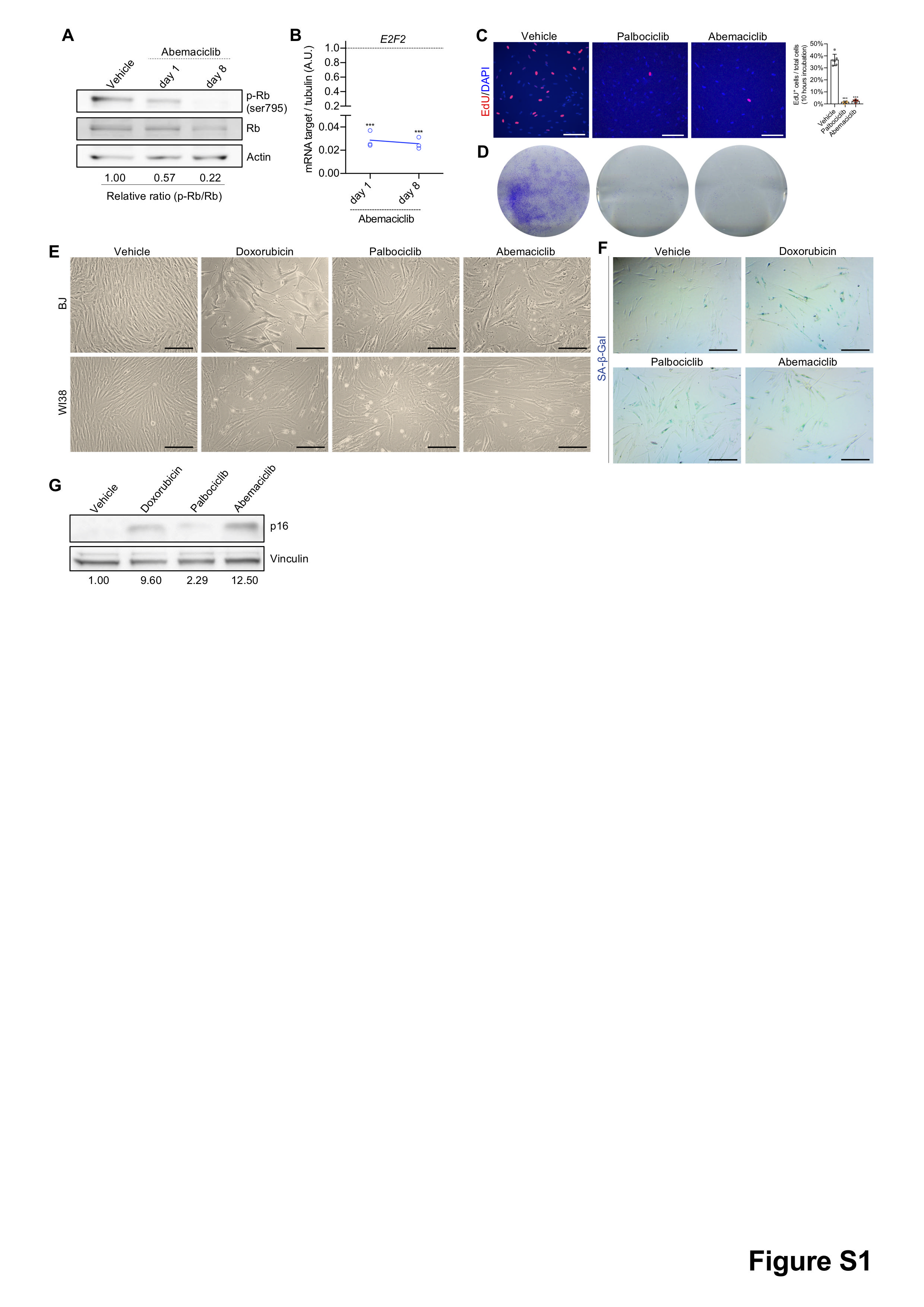
**Figure S1. CDK4/6i induces cellular senescence.**

Human fibroblasts (BJ or WI38) were treated with vehicle (water for 8 times 24 hours) or abemaciclib (1 μM for 1 or 8 times 24 hours) or palbociclib (8 times 24 hours) or doxorubicin (250 nM for 24 hours). (**A**) Protein was isolated from vehicle or abemaciclib-treated BJ fibroblasts and immunoblotted for p-Rb, Rb and actin. (**B**) RNA was isolated from treated BJ fibroblasts and mRNA levels of *E2F2* gene quantified by qPCR relative to tubulin (internal control). (**C**) At 8 dpt, treated WI38 fibroblasts were incubated with EdU for 10 hours and stained (scale bar, 150 μm; N=6). (**D**) 3*10^3^ treated WI38 cells were re-plated in 6-well dish, cultured for 8 days and stained with 0.2% crystal violet (N=3). (**E**) Representative phase contrast images of BJ or WI38 firbroblasts at the end of each drug treatment (scale bar, 1 mm; N=3). (**F**) At 8 dpt, treated WI38 fibroblasts were fixed and stained for SA-β-gal and quantified (scale bar, 1 mm; N=3). (**G**) Whole cell lysate of treated BJ fibroblats was used to immunoblot for p16 (N=3). Two-way ANOVA, data are means ±SD (**B**). One-way ANOVA, data are means ±SD (**C**). *p<0.05, ***p<0.001, N.S.=not significant. dpt, days post treatment.

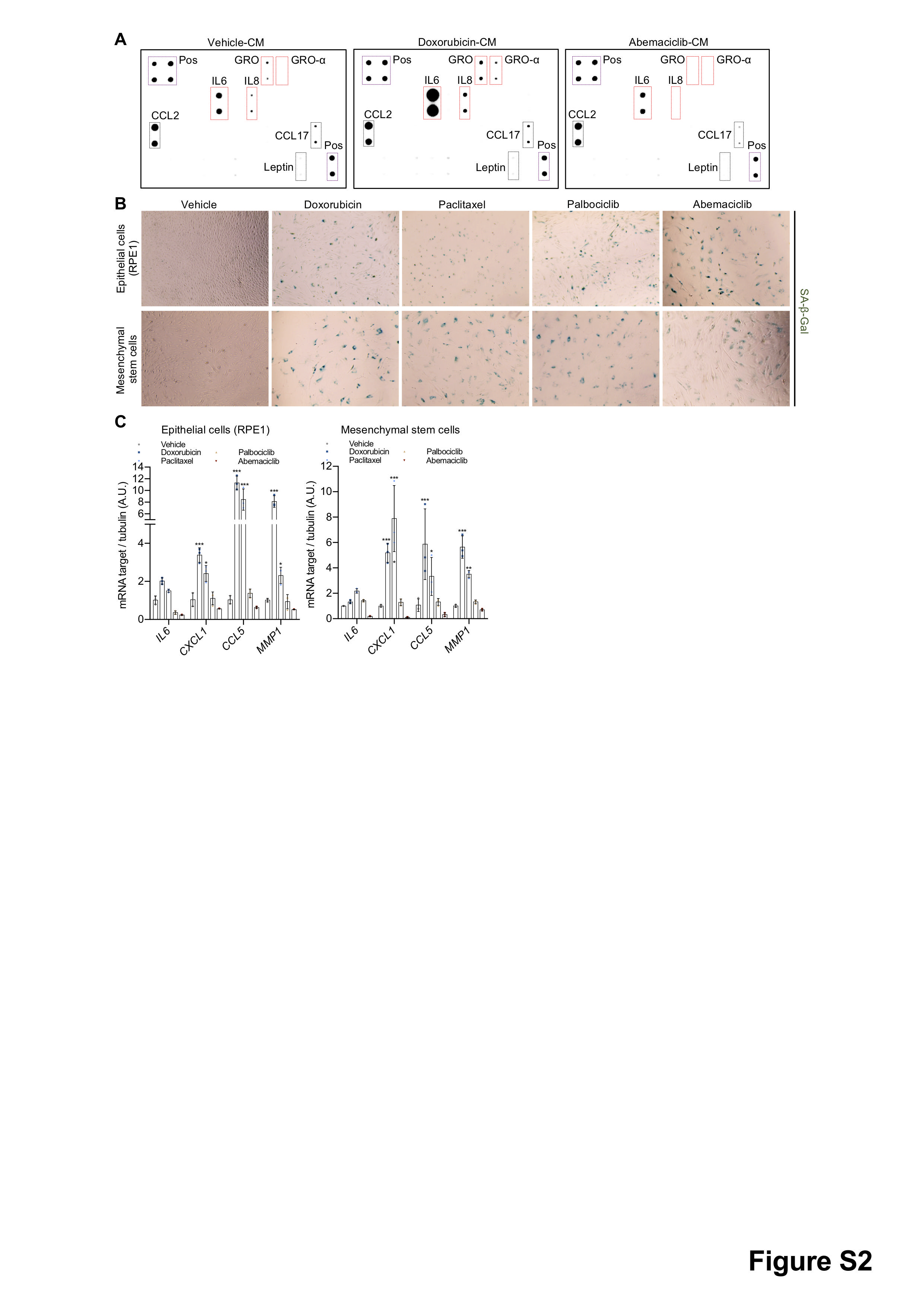

**Figure S2. CDK4/6i induces cellular senescence without NF-κB-associated SASP.**

(**A**) Proteins in the conditioned media (CM) collected from doxorubicin-(250 nM for 24 hours) or abemaciclib-(1 μM for 8 times 24 hours) treated BJ fibroblasts (8 dpt) were measured by cytokine array. (**B**) Human hTERT-RPE1 and lung mesenchymal stem cells were treated with vehicle (DMSO for 8 times 24 hours) or doxorubicin (250 nM for 24 hours) or paclitaxel (50 nM for 24 hours) or palbociclib (1 μM for 8 times 24 hours) or abemaciclib (1 μM for 8 times 24 hours). At 8 dpt, treated cells were fixed and stained for SA-β-gal (scale bar, 1 mm; N=3). (**C**) RNA was isolated from treated hTERT-RPE1 and lung mesenchymal stem cells and mRNA for the indicated NF-κB target SASP genes quantified by qRT-PCR relative to tubulin (N=3). Two-way ANOVA, data are means ±SD (**C**). *p<0.05, **p<0.01, ***p<0.001. dpt, days post treatment.

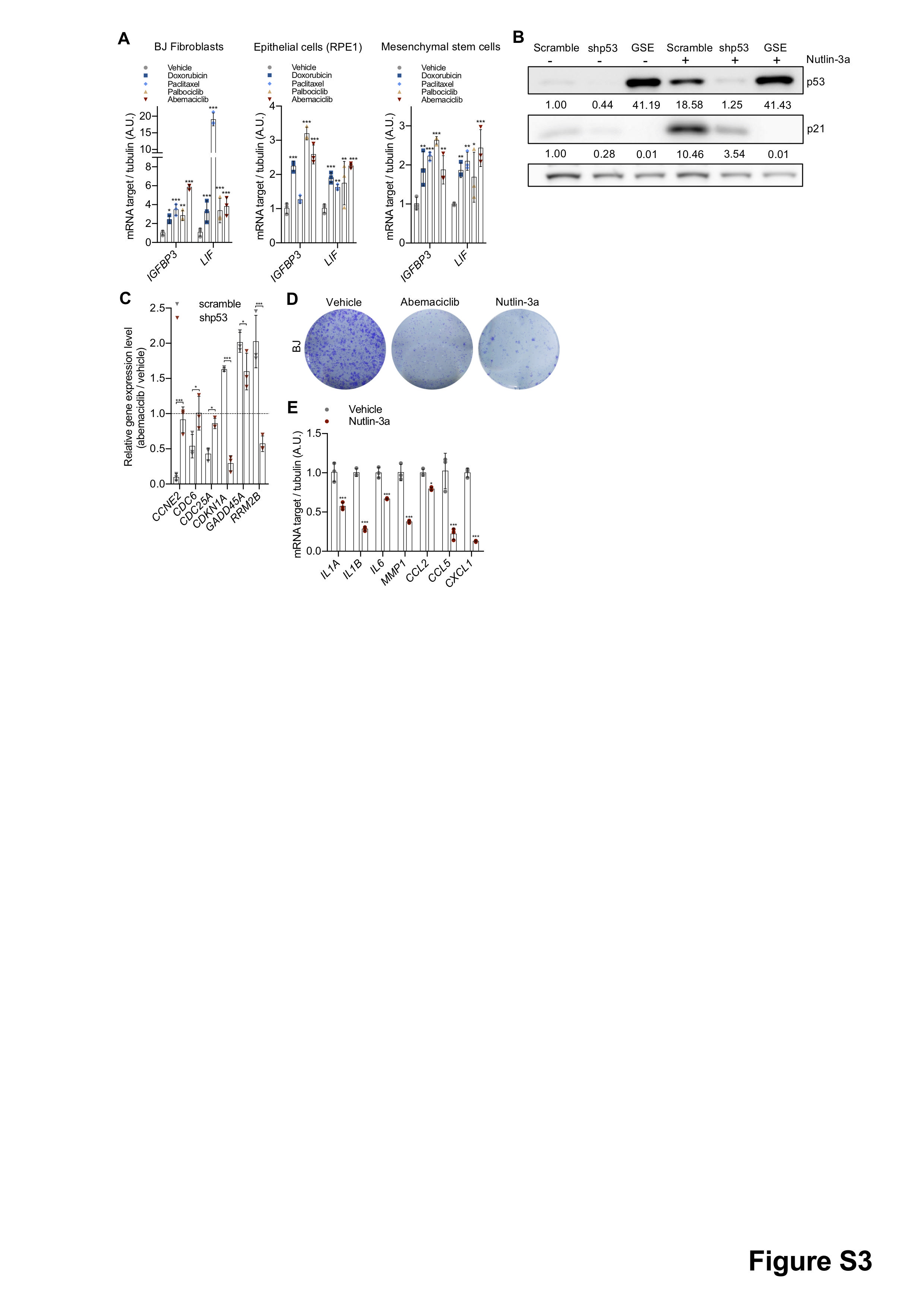

**Figure S3. CDK4/6i induces p53-associated SASP.**

(**A**) Human BJ fibroblasts, hTERT-RPE1 and lung mesenchymal stem cells were treated with vehicle (DMSO for 8 times 24 hours) or doxorubicin (250 nM for 24 hours) or paclitaxel (50 nM for 24 hours) or palbociclib (1 μM for 8 times 24 hours) or abemaciclib (1 μM for 8 times 24 hours). At 8 dpt, RNA was isolated and mRNA for the indicated p53 target SASP genes quantified by qRT-PCR relative to tubulin (N=3). (**B**) Immunoblot of p53 and p21 on BJ fibroblasts of indicated genotypes and treatments (nutlin-3a, 10 μM). GSE, gene suppressive elements of p53. (**C**) RNA was isolated from vehicle or abemaciclib (1 μM) treated scramble/shp53 BJ fibroblasts and mRNA for the indicated p53 target genes quantified by qRT-PCR relative to tubulin (N=3). (**D**) BJ fibroblasts were re-plated after treatment with vehicle (DMSO) or abemaciclib (1 μM for 8 times 24 hours) or nutlin-3a (10 μM for 8 times 24 hours) and stained with 0.2% crystal violet 8 dpt (N=3). (**E**) RNA was isolated from vehicle or nutlin-3a treated BJ cells at 8 dpt and quantified by qRT-PCR for NF-κB target SASP genes (N=3). Two-way ANOVA, data are means ±SD (**A**, **C** and **E**). *p<0.05, **p<0.01, ***p<0.001. dpt, days post treatment.

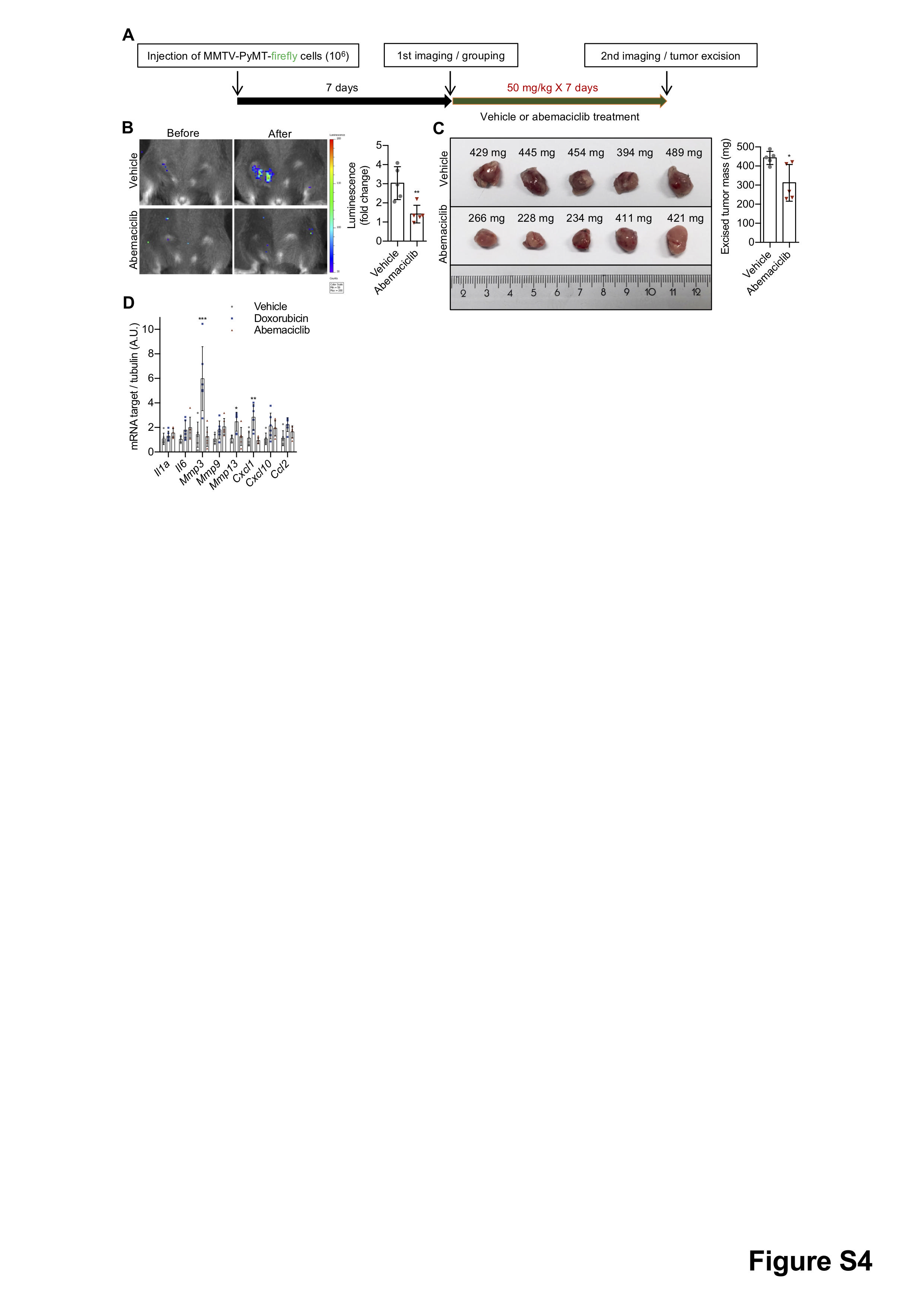

**Figure S4. Senescence-inducing dose of abemaciclib inhibits tumor growth.**

(**A**) Scheme of abemaciclib treatments for MMTV-PyMT-firefly breast cancer mouse model *in vivo*. (**B**) Female p16-3MR mice bearing MMTV-PyMT-firefly tumors in mammary fat pad were treated with vehicle (PBS, 7 consecutive days) or abemaciclib (50 mg/kg in PBS, 7 consecutive days). The mice were injected with D-Luciferin and bioluminescence was visualized/quantified by the IVIS spectrum *in vivo* imaging system before and after abemaciclib treatments, as shown by representative images and quantification (N=5 mice/group). (**C**) Excised tumors and quantification of tumor weights (N=5 mice/group). (**D**) RNA was isolated from livers of drug-treated p16-3MR mice at 15 dpt and quantified by qRT-PCR for mRNA encoding NF-κB-associated SASP genes (N=6 mice). Unpaired two-tailed *t*-test, data are means ± SD (**B** and **C**). Two-way ANOVA, data are means ± SD (**D**). *p<0.05, **p<0.01, ***p<0.001, N.S.=not significant. dpt, days post treatment.

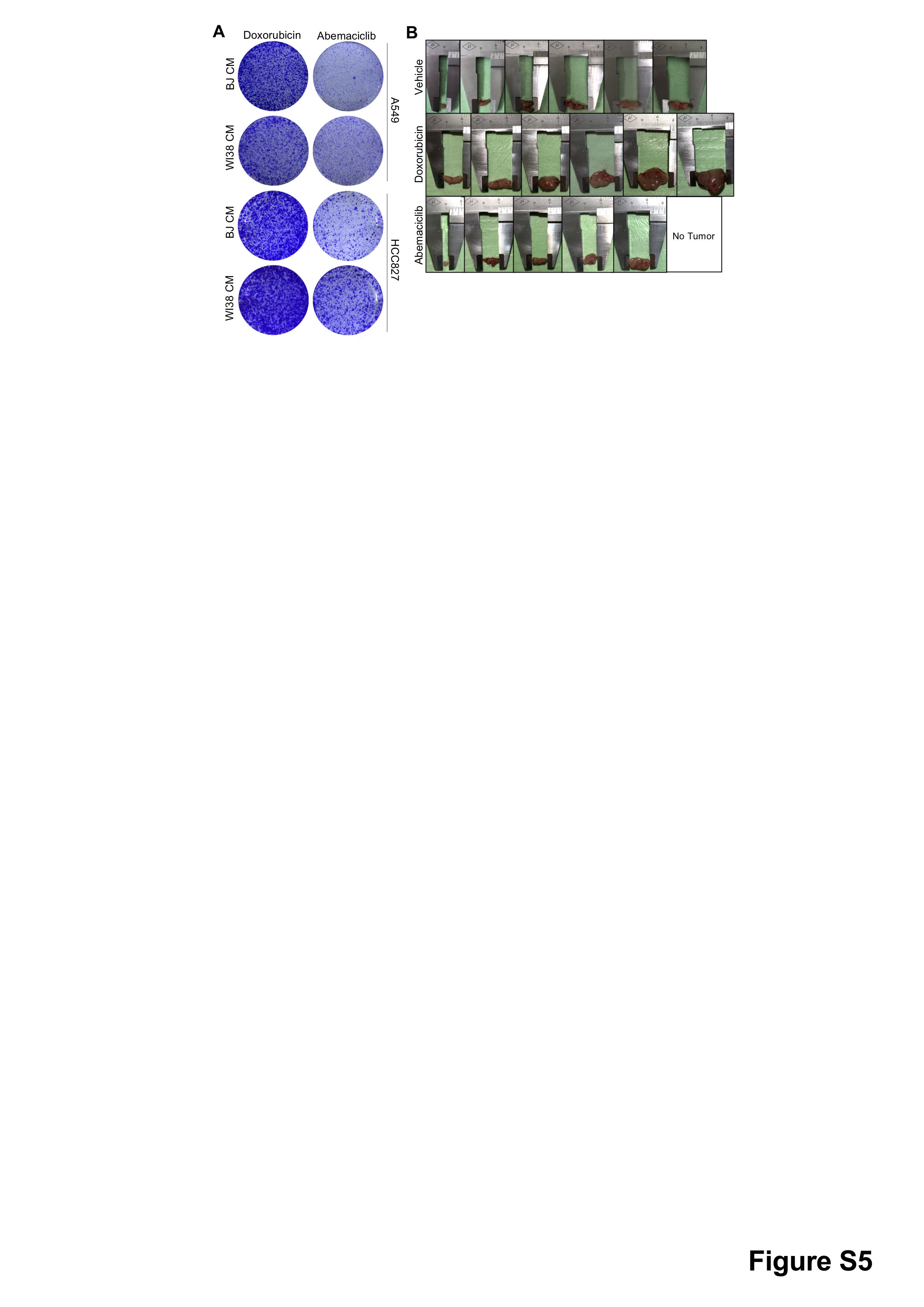

**Figure S5. p53-associated SASP lacks pro-tumorigenic properties.**

(**A**) A549 and HCC827 lung cancer cells were incubated with drug-induced BJ or WI38 serum-free CM for 45 hours. Cells were re-plated for colony formation assay for 8 days (N=3). (**B**) Excised tumors from Fig. 4B.

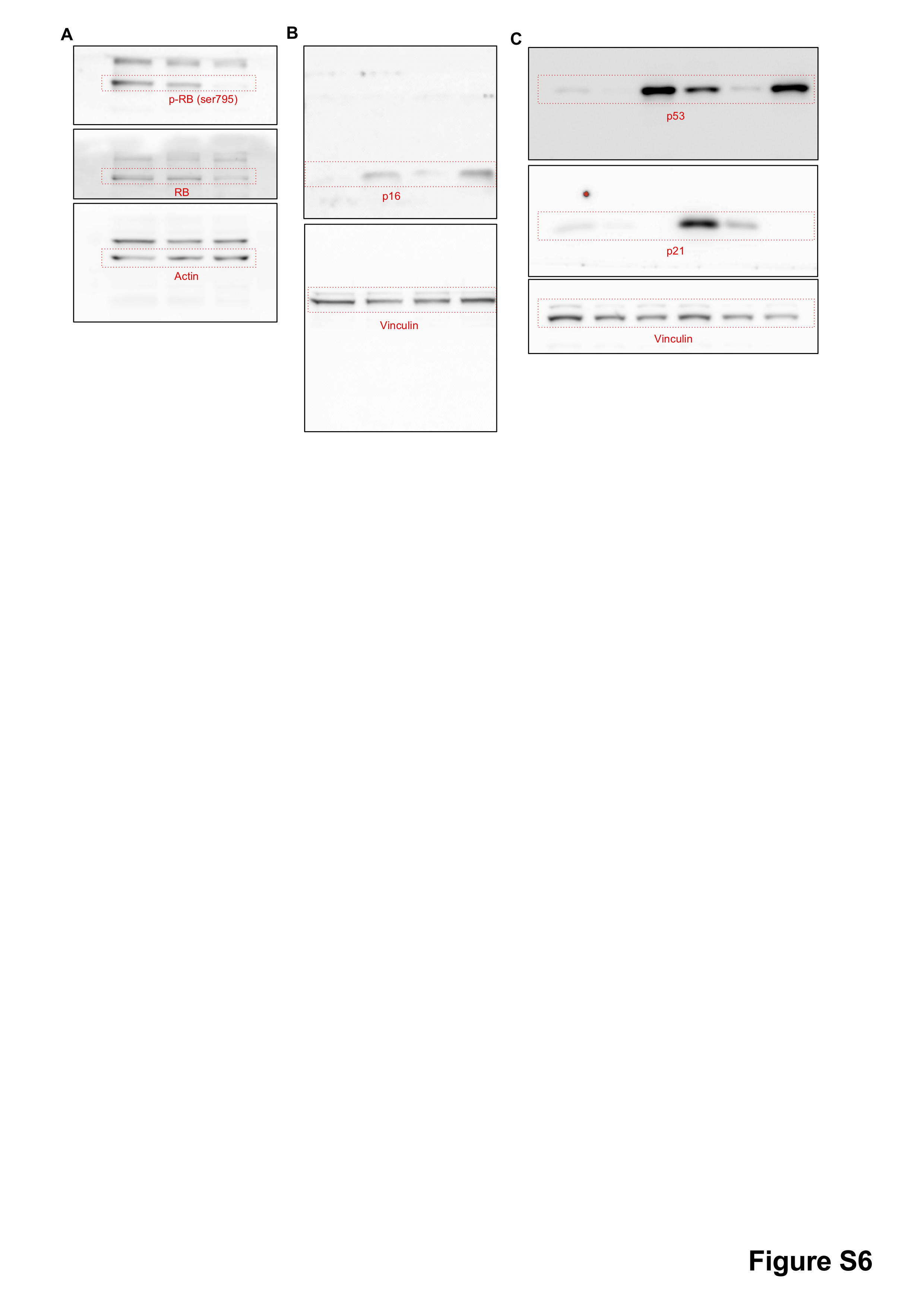

**Figure S6. Full Blots.**

(**A**) Related to fig. S1A. (**B**) Related to fig. S1G. (**C**) Related to fig. S3B.
